# AINN-P1: A Compact Sequence-Only Protein Language Model Achieves Competitive Fitness Prediction on ProteinGym

**DOI:** 10.64898/2026.03.26.714619

**Authors:** Roger Wang, Kevin Jin, Lurong Pan

**Affiliations:** Ainnocence Inc., San Francisco, CA, USA

## Abstract

Protein language models (PLMs) are increasingly central to protein engineering and drug discovery. Many high-performing systems, however, rely on large parameter counts, multiple sequence alignments (MSAs), explicit structural inputs, or computationally intensive attention mechanisms, limiting their accessibility and throughput. Here we present AINN-P1, a 167M-parameter protein language model trained exclusively on raw UniRef amino-acid sequences using an autoregressive next-token prediction objective. AINN-P1 employs a multiplicative LSTM (mLSTM) architecture—an attention-free, recurrent design that scales linearly with sequence length and avoids growing key–value caches during inference.

We evaluate AINN-P1 on ProteinGym fitness prediction tasks spanning activity, binding, expression, and stability using a frozen-encoder protocol with lightweight few-shot regression heads. Under this protocol, AINN-P1 achieves an average Spearman *ρ* of 0.441 across four task categories and a Spearman *ρ* of 0.625 on stability—the highest among sequence-only models in our comparison set. Because our evaluation uses few-shot supervised regression rather than the zero-shot scoring employed by most ProteinGym leaderboard baselines, direct numerical comparison requires caution; we discuss this methodological distinction throughout.

Beyond benchmark performance, AINN-P1 emphasizes practical deployability: its recurrent architecture avoids quadratic memory scaling, supports fixed-state inference on long sequences, and enables rapid adaptation through frozen embeddings rather than costly end-to-end fine-tuning. We discuss when sequence-only models are sufficient when structural information remains beneficial and how compact foundation models can serve as efficient front-end filters in drug discovery workflows.

## 1 Introduction

Protein engineering underpins modern biotechnology—from therapeutic antibodies and enzyme optimization to vaccine design and synthetic biology. A persistent challenge is navigating vast combinatorial sequence spaces with limited experimental budgets. Protein language models (PLMs) have reframed this problem by learning transferable representations from large sequence corpora, enabling zero-shot and few-shot prediction of mutation effects directly from amino-acid sequences.

The field has bifurcated along two axes. One axis concerns *modality*: single-sequence models versus those incorporating MSAs and/or structural information. The second concerns *scale*: from hundreds of millions of parameters to tens or hundreds of billions. While multimodality and scale have produced impressive gains, they often require training budgets, inference costs, and preprocessing pipelines (MSA search, structure prediction) that limit accessibility and throughput in applied settings.

This work explores a **sequence-first foundation model paradigm**. We ask: how far can a carefully trained, moderate-size, sequence-only model go? We present AINN-P1, a 167M-parameter model built on a multiplicative LSTM architecture and trained on UniRef sequences with an autoregressive objective. In a frozen-embedding, few-shot evaluation on ProteinGym, AINN-P1 achieves competitive performance across activity, binding, expression, and stability tasks—with particularly strong results on stability prediction.

### Contributions

- We introduce AINN-P1, a 167M-parameter sequence-only protein language model built on a multiplicative LSTM architecture, trained with an autoregressive objective on UniRef.
- We report ProteinGym fitness prediction results using a frozen-embedding, few-shot regression protocol, including strong stability performance among sequence-only models.
- We demonstrate that an attention-free, recurrent architecture achieves competitive performance while offering practical advantages in memory efficiency and inference scalability.
- We discuss practical implications for drug discovery workflows and contextualize when sequence-first models are sufficient versus when structural information adds value.

### Important methodological note

Throughout this paper, we compare AINN-P1’s few-shot frozen-embedding results against published ProteinGym leaderboard baselines. Most baselines use zero-shot scoring or task-specific evaluation protocols. Our few-shot protocol provides additional labeled supervision, which may inflate or deflate performance depending on the task and data regime. We flag this distinction wherever comparisons are made and encourage readers to interpret results accordingly.

## 2 Background and Related Work

### 2.1 Protein language models and transfer learning

PLMs adapt ideas from natural language processing by training neural networks on large protein sequence corpora using self-supervised objectives such as masked language modeling (MLM) or autoregressive prediction. Learned representations transfer to mutation effect prediction, structure prediction, and functional annotation. Families such as ESM [Rives et al., 2021] and ProGen [Madani et al., 2023] have demonstrated that single-sequence models capture evolutionary constraints sufficient for many downstream tasks without explicit structure or alignments.

### 2.2 ProteinGym benchmark

ProteinGym [Notin et al., 2024] consolidates deep mutational scanning (DMS) assays spanning multiple proteins and measurement modalities into a standardized benchmark for mutation effect prediction. Performance is primarily reported via Spearman rank correlation (*ρ*), measuring agreement between predicted and experimental fitness across mutant libraries. The ProteinGym leaderboard reports results under varying evaluation protocols (zero-shot, supervised, MSA-augmented), making cross-method comparison nontrivial.

### 2.3 Beyond sequence: MSAs and structure

Many methods incorporate MSAs to expose evolutionary co-variation signals, and others incorporate structural information to better model folding, stability, and binding. Recent multimodal approaches jointly learn from sequence and structure [Li et al., 2024]. These methods can improve accuracy—particularly for tasks where 3D context is crucial—but they introduce additional compute and data dependencies.

### 2.4 Scale and practicality

Scaling parameter counts to billions or more [Chen et al., 2024] can deliver accuracy improvements but may be prohibitive for training and deployment. A key question for applied biotechnology is whether moderate-size models can achieve high practical utility when paired with efficient adaptation protocols.

## 3 Model: AINN-P1

### 3.1 Sequence-first design philosophy

AINN-P1 is a sequence-only protein language model designed to learn transferable representations directly from raw amino-acid sequences. The model operates without MSAs, predicted structures, or external annotations at either training or inference time. Each protein is represented as a single linear sequence of residues, enabling efficient scaling and deployment in high-throughput protein engineering workflows. All biological signals are inferred implicitly from sequence statistics learned during self-supervised pretraining.

### 3.2 Architecture

AINN-P1 is built on a multiplicative Long Short-Term Memory (mLSTM) architecture [Krause et al.,2016]. Protein sequences are tokenized at the amino-acid level using a fixed vocabulary of 20 standard residues, augmented with explicit start-of-sequence and end-of-sequence tokens. Each token is mapped into a learned embedding space.

Embedded sequences are processed by a multiplicative LSTM encoder. Unlike a standard LSTM, the mLSTM introduces multiplicative interactions between hidden states within its gating mechanism, providing input-conditioned recurrent dynamics that increase modeling capacity for nonlinear residue dependencies. This architecture captures both local sequence motifs and long-range constraints arising from protein evolution.

The model uses packed sequence processing for efficient training on variable-length sequences, restricting computation to valid residues and avoiding unnecessary padding. At each sequence position, the mLSTM produces a hidden representation projected through a linear output layer to produce logits over the amino-acid vocabulary.

AINN-P1’s key architectural properties:

- **Compact footprint (167M parameters):** Enables training and deployment under constrained compute budgets.
- **Linear scaling with sequence length:** Avoids the quadratic memory cost of dense attention [Yaswani et al., 2017], which becomes a bottleneck for long protein sequences unless specialized kernels are used.
- **Fixed-state inference:** Recurrence provides constant-memory inference without the growing key-value cache footprint of attention-based decoders [Pope et al., 2023].

Figure 1 illustrates the sequence-first pipeline.

**Figure 1:**
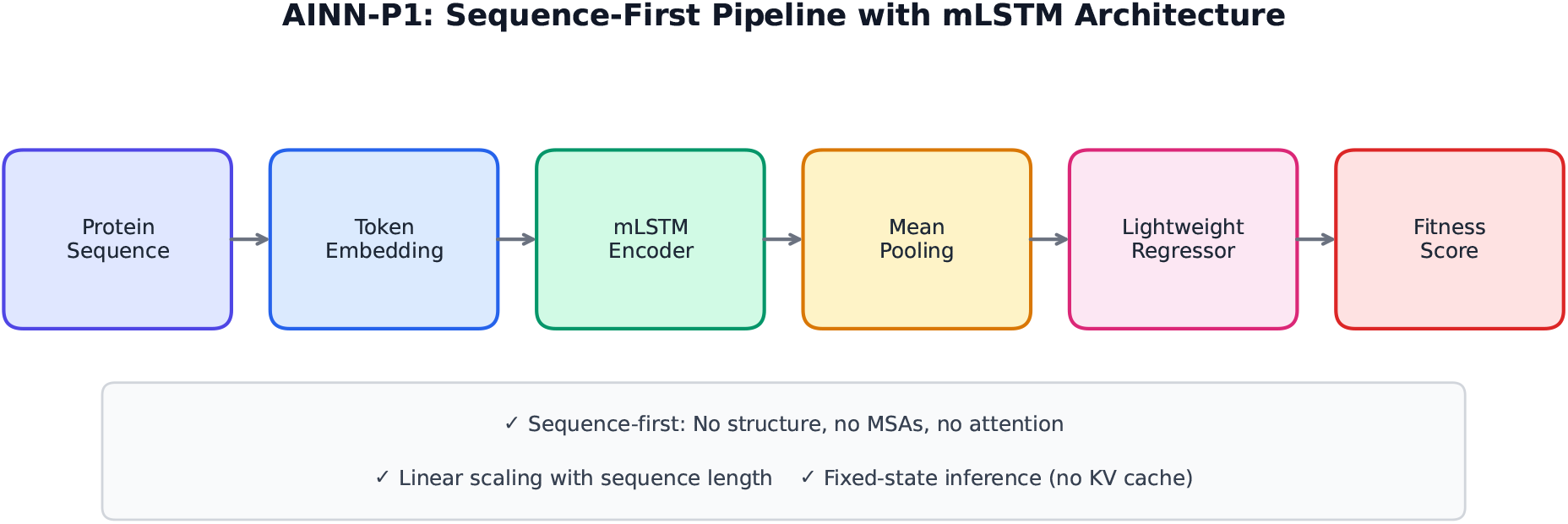
AINN-P1 sequence-first pipeline. The model is pretrained on UniRef sequences using an autoregressive objective, then used as a frozen encoder. Residue-level hidden states are mean-pooled to produce fixed-dimensional embeddings, which serve as inputs to lightweight regressors trained in a few-shot manner for each assay. This protocol avoids expensive end-to-end fine-tuning.

### 3.3 Training data and objective

AINN-P1 is trained on protein sequences from UniRef using an autoregressive next-token prediction objective. Given a protein sequence (*x*_1_, *x*_2_, …, *x*_*T*_), the model maximizes:

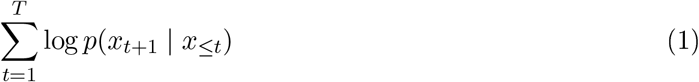

where explicit start-of-sequence and end-of-sequence tokens define sequence boundaries. Cross-entropy loss is computed at every residue position.

This causal language modeling objective encourages the model to learn both short-range residue patterns and long-range dependencies reflecting evolutionary and biophysical constraints. Unlike masked language modeling, autoregressive prediction relies solely on left-to-right context. While this means the model lacks explicit bidirectional conditioning during pretraining, the recurrent architecture naturally accumulates long-range context through its hidden state, and the approach aligns well with the mLSTM’s sequential processing.

### 3.4 Embedding extraction for downstream tasks

For downstream evaluation, AINN-P1 is used as a frozen encoder. Residue-level hidden states from the mLSTM are aggregated via mean pooling (excluding padding positions) to produce a fixed-dimensional embedding per protein sequence. These embeddings serve as inputs to lightweight regression models, enabling rapid adaptation without end-to-end fine-tuning.

## 4 Evaluation Protocol

### 4.1 Tasks

We evaluate across four ProteinGym fitness categories: **Activity** (enzymatic function), **Binding** (affinity/interaction strength), **Expression** (yield/solubility), and **Stability** (thermostability/developability). These categories span core protein engineering objectives from enzymatic function through target engagement, manufacturability, and developability.

### 4.2 Metric

For each assay, we compute Spearman rank correlation (*ρ*) between model-predicted fitness and measured experimental fitness. We report per-category averages and an overall mean across all four categories.

### 4.3 Few-shot frozen-embedding regression

To reflect realistic low-data regimes common in early-stage drug discovery, we evaluate AINN-P1 using frozen embeddings and a lightweight regressor. For each assay, we: (i) compute AINN-P1 embeddings for all mutant sequences using the frozen encoder, (ii) fit a small regressor (ridge regression) on a limited labeled subset, and (iii) report Spearman *ρ* on held-out mutants.

#### Protocol differences from leaderboard baselines

Most ProteinGym leaderboard models report zero-shot performance, where fitness scores are derived directly from model likelihoods without any labeled training data. Our protocol uses a small number of labeled examples per assay, introducing a form of supervised signal not present in zero-shot evaluation. This means our results are *not directly comparable* to zero-shot baselines—the few-shot setting may confer an advantage when labeled data is informative, or a disadvantage if the labeled subset is noisy or unrepresentative. We present both protocols’ results side by side for context, but caution against interpreting small numerical differences as definitive performance gaps.

## 5 Results

### 5.1 ProteinGym performance

Table 1 summarizes AINN-P1’s few-shot results alongside representative ProteinGym leaderboard baselines. We emphasize that baseline numbers reflect their originally reported evaluation protocols (typically zero-shot), while AINN-P1 numbers use the few-shot frozen-embedding protocol described in Section 4.3.

**Table 1:** ProteinGym comparison across four fitness categories (Spearman *ρ*). Baseline values are from published ProteinGym leaderboard results using their native evaluation protocols (typically zero-shot scoring). AINN-P1 values use few-shot frozen-embedding regression. **Bold** indicates the highest value per column; <u>underline</u> indicates the second highest. Due to protocol differences, direct numerical comparison should be interpreted with caution (see Section 4.3).

| Model (modality) | Avg $\rho$ | Activity | Binding | Expression | Stability |
| --- | --- | --- | --- | --- | --- |
| AINN-P1 (167M, seq)<sup>‡</sup> | **0.441** | 0.358 | <u>0.390</u> | 0.391 | **0.625** |
| ProSST (seq+struct) | <u>0.459</u> | <u>0.392</u> | **0.410** | **0.447** | <u>0.589</u> |
| ESM2 (150M, seq) | 0.407 | 0.391 | 0.326 | <u>0.402</u> | 0.510 |
| ProGen2-M (seq) | 0.379 | **0.393** | 0.295 | 0.433 | 0.396 |
| xTrimoPGLM-1B (seq) | 0.385 | 0.390 | 0.298 | 0.406 | 0.445 |
| xTrimoPGLM-100B (seq) | 0.366 | 0.378 | 0.259 | 0.380 | 0.450 |

<sup>‡</sup>Few-shot frozen-embedding protocol; all other rows use their published (typically zero-shot) protocols.

### 5.2 Key observations

#### Stability performance

AINN-P1 achieves a stability Spearman *ρ* of 0. 625, the highest among sequence-only models in this comparison set and competitive with the structure-augmented ProSST model (0.589). Stability is a critical proxy for developability and manufacturability in biologics, making this result practically relevant even if protocol differences complicate direct comparison.

#### Binding prediction

AINN-P1 achieves a binding *ρ* of 0. 390, substantially above similarly sized sequence-only baselines (ESM2-150M: 0. 326 ; ProGen2-M: 0.295). This suggests that sequence-only pretraining captures interaction-relevant signals that partially compensate for the absence of explicit structural inputs, though the few-shot supervision may also contribute.

#### Overall average

AINN-P1’s average *ρ* of 0.441 across all four categories is competitive, though we note that ProSST achieves 0.459 with the benefit of structural features under its native protocol. The gap narrows considerably relative to sequence-only baselines of much larger scale (xTrimoPGLM-100B: 0.366 with 600*×* more parameters).

### 5.3 Extended comparison

Table 2 provides additional context by including multimodal and larger baselines. The same protocol caveats apply.

**Table 2:** Extended ProteinGym comparison including multimodal and larger baselines (Spearman *ρ*). All baseline values are from leaderboard-style summaries. AINN-P1 uses few-shot frozen-embedding regression; comparability caveats from Table 1 apply.

| Model (modality) | Avg $\rho$ | Activity | Binding | Expression | Stability |
| --- | --- | --- | --- | --- | --- |
| AINN-P1 (167M, seq)<sup>‡</sup> | 0.436 | 0.352 | 0.394 | 0.359 | **0.637** |
| ESM3 open (multimodal) | **0.466** | **0.430** | 0.400 | **0.470** | 0.641 |
| ProSST (seq+struct) | 0.438 | 0.392 | **0.410** | 0.447 | 0.589 |
| ESM-C (300M, seq) | 0.406 | 0.423 | 0.315 | 0.408 | 0.526 |
| ESM2 (650M, seq) | 0.387 | 0.425 | 0.337 | 0.415 | 0.513 |
| xTrimoPGLM-3B (seq) | 0.399 | 0.408 | 0.308 | 0.396 | 0.482 |

<sup>‡</sup>Few-shot frozen-embedding protocol; all other rows use their published protocols.

### 5.4 Visual comparison

Figure 2 shows per-task ProteinGym performance across models, highlighting AINN-P1’s strong stability result and competitive performance across evaluation categories.

**Figure 2:**
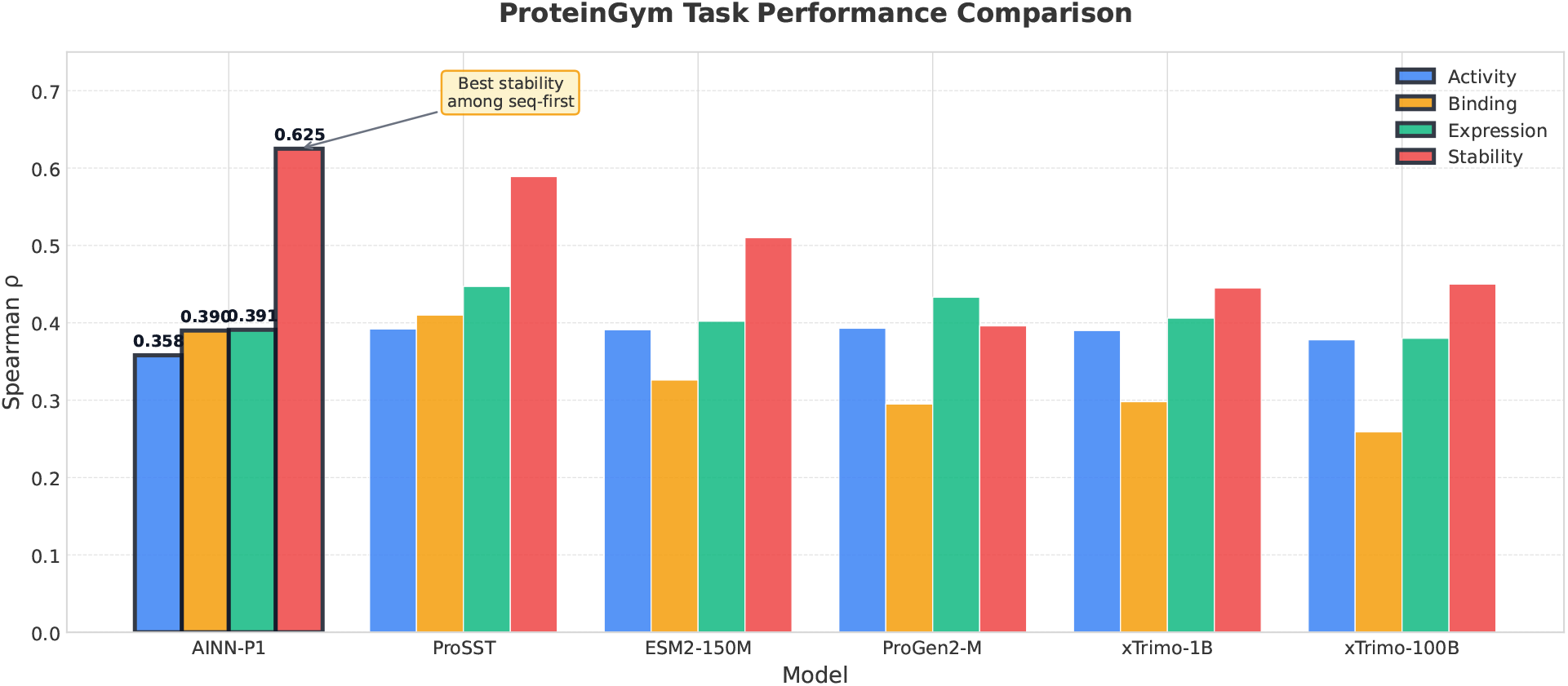
Per-task ProteinGym performance comparison. AINN-P1 is evaluated with few-shot frozen embeddings; other models reflect their published leaderboard baselines (typically zero-shot). AINN-P1 achieves the strongest stability performance among sequence-only models in this comparison set.

## 6 Discussion

### 6.1 Why sequence-first models capture stability

Why can a sequence-only model achieve strong stability prediction? A plausible explanation is that evolution compresses structural constraints into sequence distributions. Self-supervised training on diverse sequences exposes the model to residue patterns correlated with fold stability, surface exposure, and functional motifs. Recurrent architectures represent long-range dependencies that often correspond to spatial proximity in folded structures, thereby approximating structural reasoning without explicit coordinates.

Stability often depends on broad sequence statistics—hydrophobic packing, charge balance, proline/glycine placement—that are detectable from sequence context across many protein families. These “global” sequence features make stability particularly amenable to sequence-first transfer learning.

### 6.2 Binding: partial capture from sequence

Binding signals may concentrate at interfaces and depend on conformational state, creating advantages for structure-aware methods. Nevertheless, AINN-P1’s improved binding performance relative to sequence-only baselines suggests that autoregressive pretraining captures interaction motifs and compatibility constraints that generalize across proteins. We note, however, that the few-shot supervision likely contributes to this improvement, and a fair comparison would require evaluating all models under the same few-shot protocol.

### 6.3 When structure is still needed

A sequence-first model does not replace structural reasoning. Structure remains essential when function depends on multi-state conformational changes, binding requires precise geometric complementarity, engineering involves explicit steric constraints, or uncertainty is high and structure provides a mechanistic prior. In these regimes, hybrid pipelines—sequence-first screening followed by structure-aware refinement-offer a practical path forward.

### 6.4 Implications for drug discovery workflows

Sequence-first models are most impactful in platform settings where labeled data are scarce, candidate spaces are vast, and developability matters as much as peak potency. Their primary advantage is throughput: rapidly scoring large variant libraries to prioritize limited experimental budgets. In practice, they serve as a front-end triage layer in stage-gated workflows.

Concrete applications include:

- **Antibody engineering:** Early developability screens emphasizing stability and expression, complementing binding-focused methods [Jain et al., 2017, Raybould et al., 2019, Zhang et al., 2022, Bailly et al., 2020]. Sequence-only language models have also shown efficient mutation proposal without antigen structure [Hie et al., 2024], and antibody-specific resources further support domain-specialized modeling [Olsen et al., 2022a,b]
- **Small-molecule and PROTAC programs:** Prioritizing target constructs and mutation panels likely to yield robust binding and functional readouts, complementing later structure-intensive steps [Békés et al., 2022, Paiva and Crews, 2019].
- **Cell therapy optimization:** Rapid priors for ranking combinatorial receptor designs before functional testing [Mei et al., 2024], with sequence-based TCR modeling illustrating both the promise and the need for careful validation [Croce et al., 2024, Karthikeyan et al., 2025].

The primary impact is reduced wet-lab iteration and improved experiment selection rather than marginal benchmark gains. Sequence-first models excel here because they are inexpensive to deploy, adapt easily via frozen embeddings and lightweight heads, and scale as efficient first-pass filters before more complex structure-aware methods [Biswas et al., 2021, Yang et al., 2025].

## 7 Practical Guidance for Practitioners

### Choice of embedding

For global properties (stability, expression), per-sequence mean-pooled embeddings are generally adequate. For localized properties (binding interfaces, catalytic residues), per-residue embeddings pooled over annotated regions can improve sensitivity when region boundaries are known.

### Regressor selection

In low-data regimes, linear models (ridge regression, elastic net) frequently outperform more expressive regressors because they resist overfitting. With more labeled data, shallow MLPs can capture nonlinearities while remaining inexpensive. These regressors train in seconds to minutes on standard hardware.

### Variance and confidence

Few-shot performance varies across random labeled subsets. We recommend reporting confidence intervals over multiple random draws and using ensembles of small regressors for high-stakes operational decisions.

### Ranking versus calibration

Spearman *ρ* aligns with prioritization in screening pipelines. For calibrated probability estimates (e.g., will this variant pass a developability gate?), post-hoc calibration on a small validation set is advisable.

### Hybrid pipelines

A sequence-first model can serve as the fast first stage in a cascade: (i) screen and rank large libraries via embeddings + regressor, (ii) refine top candidates via structure-aware scoring, and (iii) allocate wet-lab experiments to the highest expected-value region of sequence space.

## 8 Limitations

We note several important limitations:

### Protocol comparability

The most significant caveat is that AINN-P1 is evaluated under a few-shot supervised protocol, while most baselines use zero-shot scoring. This means our results are not directly comparable to leaderboard values. A controlled comparison would require evaluating all models under the same protocol—either all zero-shot or all few-shot—which we have not done here. We plan to release zero-shot results in a future update.

### Architectural details

We do not disclose all architectural hyperparameters (hidden dimension, number of layers, learning rate schedule, training duration). This limits independent reproducibility. We plan to release model weights and training details in a future version.

### Unidirectional pretraining

As an autoregressive model, AINN-P1 processes sequences left-to-right during pretraining, lacking explicit bidirectional context. This may limit performance on tasks dominated by symmetric or highly local bidirectional interactions, though global properties like stability appear less affected.

### Domain gaps

AINN-P1 is pretrained on UniRef, which may not fully represent specialized therapeutic datasets (e.g., engineered antibodies, synthetic constructs). Performance on out-of-distribution targets should be validated experimentally.

### Single evaluation snapshot

ProteinGym evolves as new assays and models are added. Our results represent a snapshot and may not reflect the latest leaderboard state.

## 9 Conclusion

AINN-P1 demonstrates that a compact, sequence-only protein language model built on a recurrent, attention-free architecture can achieve competitive ProteinGym performance—particularly on stability, a property of high practical importance for protein engineering. The most actionable gains emerge when prediction accuracy is coupled with compute efficiency: frozen embeddings with few-shot regression provide a fast, low-cost path to useful predictions in real discovery pipelines. We believe compact sequence-first foundation models will remain a valuable component of broadly deployable protein AI systems, particularly as efficient triage layers in hybrid computational-experimental workflows.

### Reproducibility Checklist

For transparency, we document the following protocol details:

- **Assay splits:** Train/test mutants are separated by mutation identity; no duplicate sequences across splits.
- **Shot selection:** We report results with a fixed shot count per assay; variance across random draws is not yet reported but is planned for a future version.
- **Regressor:** Ridge regression with default regularization; no extensive hyperparameter search.
- **Embedding:** Mean-pooled hidden states from the final mLSTM layer.
- **Metric:** Spearman *ρ* per assay, averaged within categories and across categories.

## Data and Code Availability

AINN-P1 is trained on publicly available UniRef protein sequences. ProteinGym benchmark data are publicly available at https://proteingym.org. Model weights and evaluation code will be made available upon publication. Inquiries regarding early access may be directed to the corresponding author.

## Ethical Considerations and Responsible Use

Foundation models for biology can accelerate discovery but may also amplify risks if deployed without safeguards. We recommend treating AINN-P1 predictions as decision support rather than ground truth, pairing model-driven proposals with uncertainty estimates and confirmatory experiments. Sequence modeling capabilities could in principle be repurposed toward harmful ends; responsible deployment should include access controls, monitoring, and alignment with institutional and regulatory guidance.

## Acknowledgements

We thank colleagues at Ainnocence for discussions and internal benchmarking support.

